# Novel small molecule targeting PgQC reduces *Porphyromonas gingivalis* virulence

**DOI:** 10.1101/2025.09.30.679452

**Authors:** Nadine Taudte, Linda Liebe, Nadine Jänckel, Daniel Ramsbeck, Stephan Schilling, Jan Potempa, Sigrun Eick, Mirko Buchholz

## Abstract

Periodontitis, a chronic inflammatory disease affecting the periodontium, is primarily driven by dysbiotic of the oral microbiome with *Porphyromonas gingivalis* as a keystone pathogen. Current therapeutic approaches rely on mechanical debridement and antimicrobials, which face limitations including antibiotic resistance and microbiome disruption. Pathoblockers represent a novel therapeutic strategy that selectively targets virulence factors without bactericidal effects, potentially reducing resistance development while preserving beneficial microbiota. Here, we describe the characterization of S-0636, a novel reversible inhibitor of zinc-dependent glutaminyl cyclase (PgQC), as a compound to selectively suppress growth of *P. gingivalis*.

The compound’s effects was assessed through enzymatic assays, bacterial growth studies, virulence factor activity measurements (gingipain activity, hemagglutination, keratinocyte invasion), selectivity testing against commensal oral bacteria, resistance development analysis over 50 passages, and cytotoxicity evaluation in human cell lines.

S-0636 demonstrated potent PgQC inhibition with a K_i_ value of 0.014 μM and has successfully reduced the intracellular PgQC activity by 50% at 8 μM and had no bactericidal effects. Treatment of *P. gingivalis* with S-0636 significantly decreased gingipain activity, impaired hemagglutination capacity, and reduced keratinocyte invasion by 76% at 62.5 μM. The compound showed high selectivity, with no growth inhibition of ten tested oral commensal species at concentrations up to 0.25 mM. Importantly, no resistance development was observed after 50 bacterial passages, and cytotoxicity remained minimal in human cell lines with >80% viability at 0.5 mM.

In previous studies, PgQC was suggested as an enzyme responsible for pGlu-modification and stabilization of bacterial virulence factors. The current study now validates PgQC as an attractive target for pathoblocker development, demonstrating that S-0636 effectively attenuates *P. gingivalis* pathogenicity through selective virulence factor inhibition while preserving bacterial viability and oral microbiome integrity. The absence of resistance development and low cytotoxicity profile support the potential clinical translation of this approach for periodontal disease management, representing a promising alternative to conventional antimicrobial therapies.

## Introduction

Periodontitis is a chronical inflammation of the periodontium, it can be caused by insufficient oral hygiene and the resulting bacterial plaque in combination with external lifestyle factors or existing underlying diseases (Kinane et al., 2017; Kozak and Pawlik, 2023). It is seen as a dysbiotic condition of the oral microbiome. The progression is characterized by a stepwise movement from the healthy “eubiotic” stage through the reversible stage of gingivitis, which can be easily managed by a consequent oral hygiene regime, and in the latest stage to the manifested periodontitis (Abdulkareem et al., 2023; Radaic and Kapila, 2021). It is one of the most prevalent diseases worldwide, in Germany every second adult (>35 years) is affected (Aimetti et al., 2015; Brauckhoff et al., 2009).

One of the most recognized pathogens involved is *P. gingivalis*, considered as a “keystone pathogen”(Hajishengallis, 2014; Hajishengallis et al., 2012). This Gram-negative bacterium expresses its virulence primarily through a range of surface-exposed and secreted proteins, particularly those embedded in the outer membrane or delivered on the surface of outer membrane vesicles (OMVs) (Lei et al., 2025; Okamura et al., 2021) For example, gingipains, *P. gingivalis* major cysteine proteases, play a central role in its pathogenicity (Lei et al., 2025). By degrading host proteins, gingipains facilitate to nutrient acquisition for both *P. gingivalis* and the surrounding microbial community. They also facilitate bacterial adherence to host cells such as keratinocytes and erythrocytes, promote the breakdown of host tissues, and interfere with immune recognition, allowing the bacterium to evade host immune defence mechanisms (Ciaston et al., 2022; Grenier and Tanabe, 2010; Socransky et al., 1998; Song et al., 2021; Zheng et al., 2021).

In addition to local inflammation, tissue destruction and bone resorption, there is growing scientific evidence linking periodontal disease to a range of systemic disorders. These associations are primarily mediated through inflammatory pathways and bacterial dissemination. The connection between periodontitis and cardiovascular disease is particularly well established. Numerous epidemiological studies consistently show that individuals with periodontal disease have an increased risk of atherosclerotic cardiovascular events, including coronary heart disease and stroke (Sanz et al., 2010, 2020a). Moreover, both epidemiological and mechanistic studies have revealed bidirectional relationships between periodontitis and several systemic disorders. For instance, diabetes mellitus not only increases the risk of developing periodontitis, but severe periodontitis can also impair glycemic control and exacerbate diabetic complications (Păunică et al., 2023; Preshaw et al., 2011; Preshaw and Bissett, 2019). Similar bidirectional links have been observed with inflammatory bowel disease, likely mediated via the oral–gut microbiome axis (Wang et al., 2024; Yamazaki, 2023; Zhou et al., 2023). Periodontitis has also been associated with respiratory diseases, rheumatoid arthritis, certain cancers (particularly oropharyngeal), and neurodegenerative disorders such as Alzheimer’s disease (Martínez-García and Hernández-Lemus, 2021). These associations are driven by chronic inflammation, microbial translocation, immune dysregulation, and pathogen-specific mechanisms, underscoring the systemic relevance of periodontal health.

The primary objective of current therapeutic strategies in periodontitis management is the control of bacterial plaque. Standard treatment involves mechanical debridement through subgingival instrumentation, including scaling and root planing (SRP), performed manually or with ultrasonic devices, often supported by pocket disinfection using chlorhexidine. In advanced cases, surgical interventions such as flap procedures may be necessary to access and clean affected areas. Adjunctive therapies may include systemic antibiotic administration, commonly using the so-called Van-Winkelhoff cocktail (amoxicillin combined with metronidazole), or local application of tetracyclines such as minocycline or doxycycline. Current S3-guidelines recommend only locally or systemically administered anti-infectives; other adjunctive therapies are generally not advised (Papapanou et al., 2018; Sanz et al., 2020b).

It is important to note that both antibiotics and disinfectants have limitations. In particular, systemic antibiotic therapy is not recommended as a first-line treatment due to increasing antibiotic resistance and a notable incidence of adverse effects, including general malaise (12.3%) and diarrhea (10.3%) (Abe et al., 2022; Heta and Robo, 2018). The continuous emergence of antibiotic-resistant pathogens poses a serious challenge to future healthcare. To break the vicious cycle of developing new antibiotics only to face subsequent resistance, a novel therapeutic strategy has been introduced: the use of compounds referred to as pathoblockers or more general next-generation antimicrobials (NGAs) (Brown and Wright, 2016; Calvert et al., 2018; Dickey et al., 2017). These agents selectively target bacterial virulence factors, reducing pathogenicity without exerting bactericidal effects. This targeted mechanism renders the microorganisms more susceptible to immune system responses and enhances the effectiveness of conventional antibiotics, while significantly lowering the risk of resistance development (Allen et al., 2014; Clatworthy et al., 2007).

Currently, a variety of strategies are being pursued to effectively reduce bacterial pathogenicity. Some small molecules are designed to inhibit bacterial adhesion, thereby preventing colonization and biofilm formation (Benz and Schmidt, 1992; Svensson et al., 2001). Others aim to disrupt quorum sensing (QS), the bacterial communication system that regulates virulence and synchronise their responses to environmental clues (Sully et al., 2014) or to interfere with intracellular signalling cascades that control key cellular processes (Sambanthamoorthy et al., 2012). A particularly promising approach involves the irreversible inhibition of the lysine-specific gingipain Kgp, which alongside the arginine-specific gingipain Rgp, represents a key virulence factor of *P. gingivalis.* This strategy is exemplified by COR588, a next-generation small-molecule inhibitor currently undergoing clinical evaluation (Raha et al., 2020).

Recently, in collaboration with research partners, we identified a protein that is essential for a range of bacteria belonging to the phylum Bacteroidetes (Hutcherson et al., 2016; Klein et al., 2012). This enzyme, bacterial glutaminyl cyclase (bacQC, PgQC), catalyzes the cyclization of N-terminal glutaminyl residues to pyroglutamate in a variety of proteins (Taudte et al., 2021). In *P. gingivalis*, it was shown that over 70% of all secreted proteins possess an N-terminal glutamine residue following the signal peptide. Many of these proteins are chaperons, cargo proteins, periplasmic proteins involved in peptide metabolism, functional or structural components of the outer membrane as well as elements of the Type IV and Type IX secretion systems (Bochtler et al., 2018; Szczęśniak et al., 2023). This highlights PgQC as an attractive target for the development of selective inhibitors aimed at compromising *in vivo* fitness of *P. gingivalis* and reducing its pathogenicity. Hence, we successfully identified the first inhibitors of this enzyme, based on a tetrahydroimidazo[4,5-c]pyridine scaffold. These compounds exhibited potent activity in the low nanomolar range, highlighting their potential as selective modulators of bacterial glutaminyl cyclase function (Ramsbeck et al., 2021).

Subsequent development led to the identification of a small molecule S-0636 with enhanced inhibitory potential, improved efficacy and selectivity, making it a promising candidate for further therapeutic exploration.

## Materials and Methods

### Synthesis of compound S-0636 (1-(2-(4 -((lmidazo[4, 5-b]pyridin-6-ylamino)methyl)-phenoxy)ethyl)guanidine)

The last step of the synthesis was caried out as follows: A solution of N-(4-(2-Aminoethoxy)benzyl)imidazo[4,5-b]pyridin-6-amine*TFA (41 mg, 0.08 mmol, 1 eq) in DMF was treated with N,N’-Bis(tert-butoxycarbonyl)-S-methylisothiourea (23 mg, 0.08 mmol, 1 eq), DMAP (1 mg, 0.008 mmol, 20 0.1 eq) and TEA (33 μI, 0.24 mmol, 3 eq). The mixture was stirred for 16 hours at room temperature, quenched with water and extracted with EtOAc (3×20 ml). The combined organic layers were dried over MgCO_3_ and evaporated. Boc-deprotection was carried out with TFA/DCM (1:1, 2 ml) and TIS (100 μI). The volatiles were evaporated, and the residue was purified by semi-preparative HPLC. Yield: 18 mg (41 %); MS m/z: 326.1 [M+H]^+^; HPLC (Gradient A): rt 5.95 min, 100%; 1H-NMR (DMSO-d6) 8: 4.04 (t, 2H, 3J=5.3 Hz); 4.30 (s, 2H); 6.91-6.95 (m, 2H); 7.07 (d, 1 H, 4J=2.5 Hz); 7.32-7.36 25 (m, 2H); 7.64 (t, 1 H, 3J=5.8 Hz); 8.09 (d, 1 H, 4J=2.4 Hz); 8.86 (s, 1 H).

### Cell culture and culture conditions

The cell lines hTERT-TIGK (human telomerase immortalized gingival keratinocytes) and hTERT-TIGF (human telomerase immortalized gingival fibroblasts) were obtained from American Type Culture Collection (ATCC,CRL-3397). hTERT-TIGK cells were cultured in Gibco™ Medium 154 supplemented with the Human Keratinocyte Growth Kit (Gibco). hTERT-TIGF cells were maintained in Dulbecco’s Modified Eagle Medium (DMEM/F12) supplemented with 10% fetal bovine serum (FBS) and GlutaMAX™ (Gibco). The HepG2 cell line was cultured in RPMI medium with 10% FBS, while HEK 293 cells were grown in DMEM supplemented with 10% FBS. All cell lines were seeded at a density of 5 × 10⁵ cells per 25 cm² and incubated at 37 °C in a 5% CO₂ atmosphere. Subculturing was performed when cells reached 70–90% confluency. For cell detachment, a 0.05% (w/v) trypsin-EDTA solution (Gibco) was used for all cell lines except TIGK, which were detached using a 0.25% (w/v) trypsin-EDTA solution. Culture medium was refreshed twice per week. Cell number and viability were assessed using trypan blue exclusion and measured with the TC20™ Automated Cell Counter (Bio-Rad).

### Bacterial strains and culture conditions

All *P. gingivalis* strains were cultured either in pre-reduced Caso Broth (Carl Roth, Germany), which is compositionally equivalent to Tryptic Soy Broth (TSB), supplemented with 5 µg/mL hemin, 1 µg/mL vitamin K₁, and 0.5 mg/mL cysteine hydrochloride, or on Caso Agar (Carl Roth, Germany) enriched with the same supplements and 5% defibrinated sheep blood (Oxoid). Cultures were incubated under anaerobic conditions at 37 °C using the GasPak™ EZ Anaerobe Gas Generating System (BD Becton Dickinson). For selectivity testing, the following bacterial strains were obtained from DSMZ: *Actinomyces oris* (DSM 43798), *Capnocytophaga granulosa* (DSM 11449), *Lautropia mirabilis* (DSM 11362), *Rothia dentocariosa* (DSM 112005), *Neisseria subflava* (DSM 17610), *Corynebacterium matruchotii* (DSM 20635) and *Haemophilus parainfluenzae* (DSM 8978). *Veillonella parvula* ATCC 17745, *Streptococcus sanguinis* ATCC 10556 and *Streptococcus mitis* ATCC 49456 were kindly provided by Prof. Sigrun Eick. All strains were cultivated according to DSMZ guidelines and additional protocols described in the Supplementary Information.

### Glutaminyl cyclase activity and inhibition assay

Recombinant *P. gingivalis* glutaminyl cyclase was expressed and purified as already described (Taudte et al., 2021). Enzymatic activity was assessed using the fluorogenic substrate H-Gln-AMC, following previously established protocols (Taudte et al., 2021). For inhibitor testing, compounds were added to the reaction mixture in the presence of 1% (v/v) DMSO. Inhibitory constants were determined by varying the substrate concentration from ¼ × K_M_ to 2 × K_M_. Reactions were initiated by the addition of 20 nM PgQC, and progress curves were fitted to the standard equation for competitive inhibition using GraFit software (Version 7, Erithacus Software Ltd., Horley, UK). For Determination of QC activity in planktonic *P. gingivalis* culture, cells were incubated with increased concentration of S-0636 for 42 h and resuspended to an OD₆₀₀ of 2. A 10 µl aliquot of the resulting suspension was assessed in the PgQC activity assay.

### Growth of *P. gingivalis* in presence of S-0636

Overnight cultures of *P. gingivalis* strains were adjusted to McFarland standard 4 in pre-reduced medium. The bacterial suspensions were then diluted 1:100 into fresh medium containing a serial 1:2 dilution of either S-0636 or minocycline dihydrate. Cultures were incubated at 37 °C under anaerobic conditions using the GasPak™ EZ Anaerobe Gas Generating System (BD Becton Dickinson). After 42 hours, bacterial growth was quantified by measuring optical density at 600 nm (OD₆₀₀) to assess the impact of the compounds. All experiments were performed in at least three independent biological replicates.

### Rgp-gingipain-activity assay

*P. gingivalis* strains were incubated with a serial 1:2 dilution of S-0636 as described above. After 42 h of growth, the cultures were centrifuged at approximately 7000 × g for 5 minutes and resuspended in PBS to an OD₆₀₀ of 2. A 10 µl aliquot of the resulting suspension was used for the Arg-gingipain activity assay, as previously described (Bochtler et al., 2018) employing the chromogenic substrate benzoyl-L-arginine-p-nitroanilide (L-BApNA, Peptanova). The reaction rate (slope/min) was determined by measuring the release of p-nitroaniline at 405 nm using a FluoStar plate reader (BMG Labtech). The assay was performed at least three times in independent experiments.

### Hemagglutination assay

*P. gingivalis* strains were incubated with a serial 1:2 dilution of S-0636, as described above. After incubation, the cultures were centrifuged and resuspended in PBS to an OD₆₀₀ of 3. Subsequently, 100 µl of the cell suspension was added to each well of a 96-well round-bottom microtiter plate and serially diluted in PBS (1:2 to 1:32). Defibrinated sheep blood (Oxoid) was washed with PBS and prepared as a 2% solution in PBS. Then, 100 µl of the sheep blood solution was added to the 100 µl of *P. gingivalis* cell suspension in each well and incubated for 3 hours at room temperature. Hemagglutination activity was assessed visually and quantified using ImageJ and GraphPad Prism 10. The assay was performed at least three times in independent experiments.

### Invasion assay

TIGKs were seeded in a technical duplicate into 24-well plates at a density of 5 × 10⁴ cells per well and cultured until reaching 80% confluency. Control wells were used for cell counting and assessment of cell viability prior to infection with *P. gingivalis* W83. In parallel, *P. gingivalis* cells were preincubated with various concentrations of S-0636 for 42 h under anaerobic conditions. Optical density was measured, and the bacteria were resuspended in Gibco™ Medium 154 to achieve a multiplicity of infection (MOI) of 200. TIGKs were infected with *P. gingivalis* for 90 minutes at 37 °C in a 5% CO₂ atmosphere, followed by antibiotic treatment for 1 hour as previously described (Ye et al., 2017). After treatment, the medium was discarded, and the cells were rinsed three times with PBS. TIGKs were lysed with 300 µl of ice-cold sterile water for 30 minutes and harvested by scraping. Serial dilutions of the lysates were plated in duplicate on blood agar plates to determine colony-forming units (CFU) of *P. gingivalis* W83. Plates were incubated for 5–7 days at 37 °C under anaerobic conditions.

### Growth response of oral commensals to S-0636 exposure

Growth experiments were conducted in 96-well plate format using defined media supplemented with increasing concentrations of S-0636. Pre-cultures were adjusted to either McFarland standard 0.5 or 4, and subsequently diluted 1:50 or 1:100 into fresh media containing a serial 1:2 dilution series of S-0636. Cultures were incubated at either 30 °C or 37 °C under anaerobic, aerobic, or 5% CO₂ conditions until reaching the stationary phase (detailed protocol available in the Supplementary Information). Optical density at 550 nm was measured using a FluoStar Optima plate reader (BMG Labtech) to assess the impact of the compound on bacterial growth. All experiments were performed in at least three independent biological replicates.

### Assessment of resistance development in *P. gingivalis* upon long-term exposure to S-0636

To investigate whether prolonged exposure to S-0636 induces resistance mechanisms in *P. gingivalis*, various strains were passaged on Caso Agar supplemented with 16 µM S-0636 for 50 passages. Cultures were maintained anaerobically at 37 °C in 6-well plates. Agar plates without compound served as controls. Passaging was performed twice weekly. After every 10 passages, the strains were cultured in Caso Broth without S-0636 for 24 hours and subsequently subjected to a dose–response experiment as described above. After 42 hours of incubation, bacterial growth and gingipain activity were assessed.

### Cytotoxicity assessment of S-0636 in human cell lines

To assess cytotoxicity, human cell lines TIGK, TIGF, HepG2, and HEK293 were seeded into 96-well plates at a density of 2 × 10⁴ cells per well in 100 µl of culture medium and incubated for approximately 24 hours at 37 °C in a 5% CO₂ atmosphere. Once the cells reached ∼80% confluency, the medium was replaced with fresh medium containing various concentrations of S-0636, followed by an additional 24-hour incubation under the same conditions. Cells treated with 0.1% Triton X-100 served as a negative control (complete cytotoxicity), while cells treated with the S-0636 compound diluent served as a positive control (no cytotoxicity). After treatment, WST-8 reagent (MCE®, MedChem Express) was added at a 1:10 dilution and incubated for 2–3 hours at 37 °C. Cell viability was quantified by measuring the absorbance of WST-8 formazan at 450 nm using a Tecan Sunrise microplate reader. All experiments were performed in at least three independent biological replicates.

## Results

### Effect of S-0636 on PgQC activity and growth of *P. gingivalis*

Recently, we published a study demonstrating the continued evolution of inhibitors targeting bacterial glutaminyl cyclase from *P. gingivalis*. Based on well-known inhibitors of the related human enzyme and a previously reported imidazo[4,5-b]pyridine derivative as the starting point, our medicinal chemistry approach has identified a novel class of tetrahydroimidazo[4,5-c]pyridine derivatives as potent inhibitors, featuring a new metal binding group that has not previously been described for glutaminyl cyclase inhibitors (Ramsbeck et al., 2021). In this paper structure–activity relationships were investigated, and the most potent inhibitor, compound 8t, demonstrated a K_i_ value of 0.084 µM. For agents acting on Gram-negative bacteria, potent inhibition alone is not sufficient; optimal compounds should also possess relatively high solubility and polarity, as shown by re-analyses of physicochemical properties of successfully marketed antibiotics (O’Shea and Moser, 2008). However, within the class of tetrahydroimidazo[4,5-c]pyridines, it was not possible to obtain compounds combining both required features. Therefore, we further investigated the already known class of imidazo[4,5-b]pyridines. Subsequent optimization of both compound properties led to the discovery of compound S-0636, which, due to its guanidinium moiety, displays sufficient polarity and solubility, while its inhibitory activity is superior to all previously known PgQC inhibitors, based on its K_i_ value of 0.014 µM, as shown in Figure 1.

**Fig. 1:**
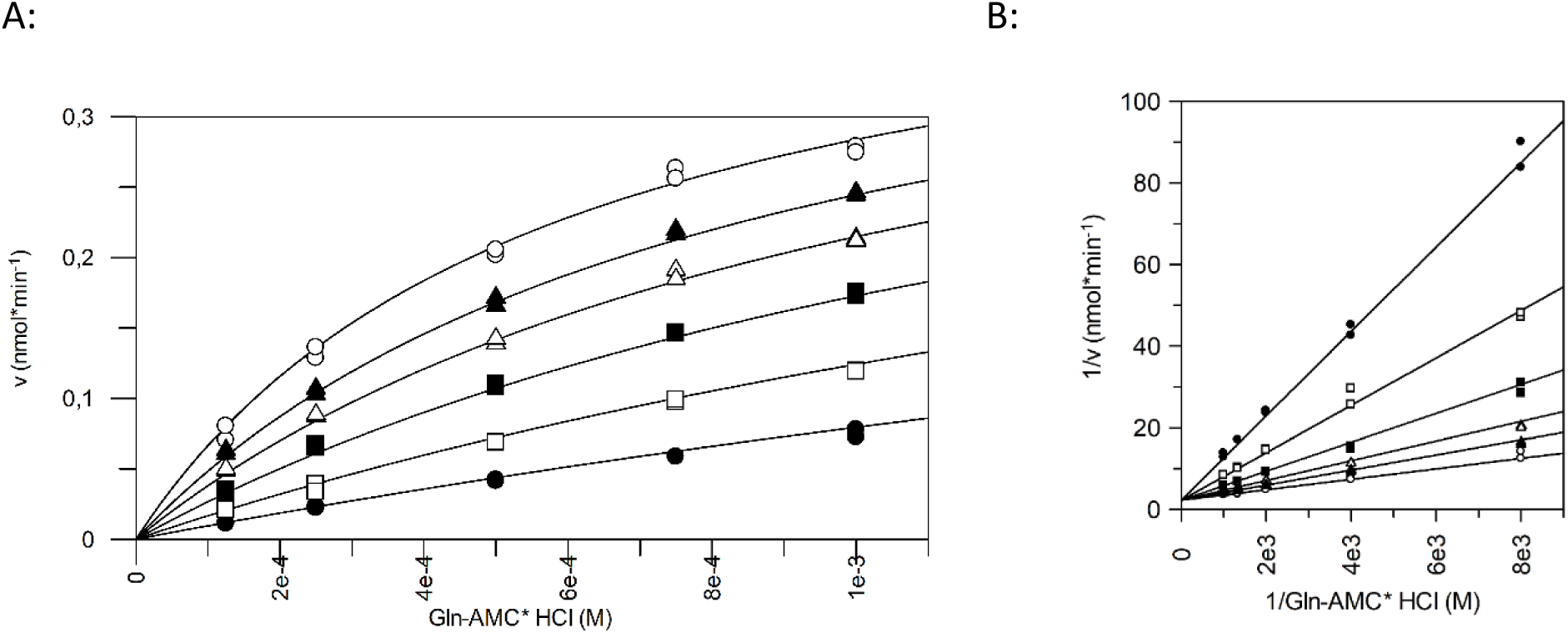
Determination of inhibitory constant of S-0636. Inhibitory constant was determined using different concentration of H-Gln-AMC varying from ¼ K_M_-2K_M_. The inhibitory constant was evaluated by fitting the set of progress curves to the general equation for competitive inhibition using GraFit software (Version 7, Erithacus software Ltd., Horley, UK), (A) exemplary v/S characteristic, (B) Lineweaver Burk of H-Gln-AMC turn over in presence of (o) w/o S-0636, (●) 100 nM, (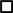) 50 nM, (■) 25 nM, (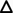) 12.5 nM and (▲) 6.25 nM in presence of 20 nM PgQC at 30°C. The inhibitory constant was performed in at least three independent technical replicates

Previous studies have shown that PgQC is localized in the periplasm, anchored to the inner membrane (Bender et al., 2019; Bochtler et al., 2018). Such subcellular localization often challenges inhibitor effectiveness, as potential compounds may not reach their target protein. As discussed above, this limitation is frequently linked to the compound’s lipophilicity, which can reduce cellular uptake and activate efflux. In contrast, S-0636 lacks significant hydrophobic properties. It was specifically designed to diffuse through the outer membrane because of its polar features and to competitively inhibit the target enzyme PgQC. To verify this mechanism, *P. gingivalis* ATCC 33277 and W83 were incubated with various concentrations of S-0636 for 42 hours, and PgQC activity was then measured in each sample. As shown in Figure 2A, PgQC activity decreased in a dose-dependent manner upon treatment with S-0636. Remarkably, even at a concentration of 8 µM, a 50% reduction in PgQC activity was observed. These data indicate that S-0636 successfully penetrates *P. gingivalis* cells and inhibits PgQC activity both *in vitro* and in live-cell assays, confirming periplasmic effectiveness.

**Fig. 2:**
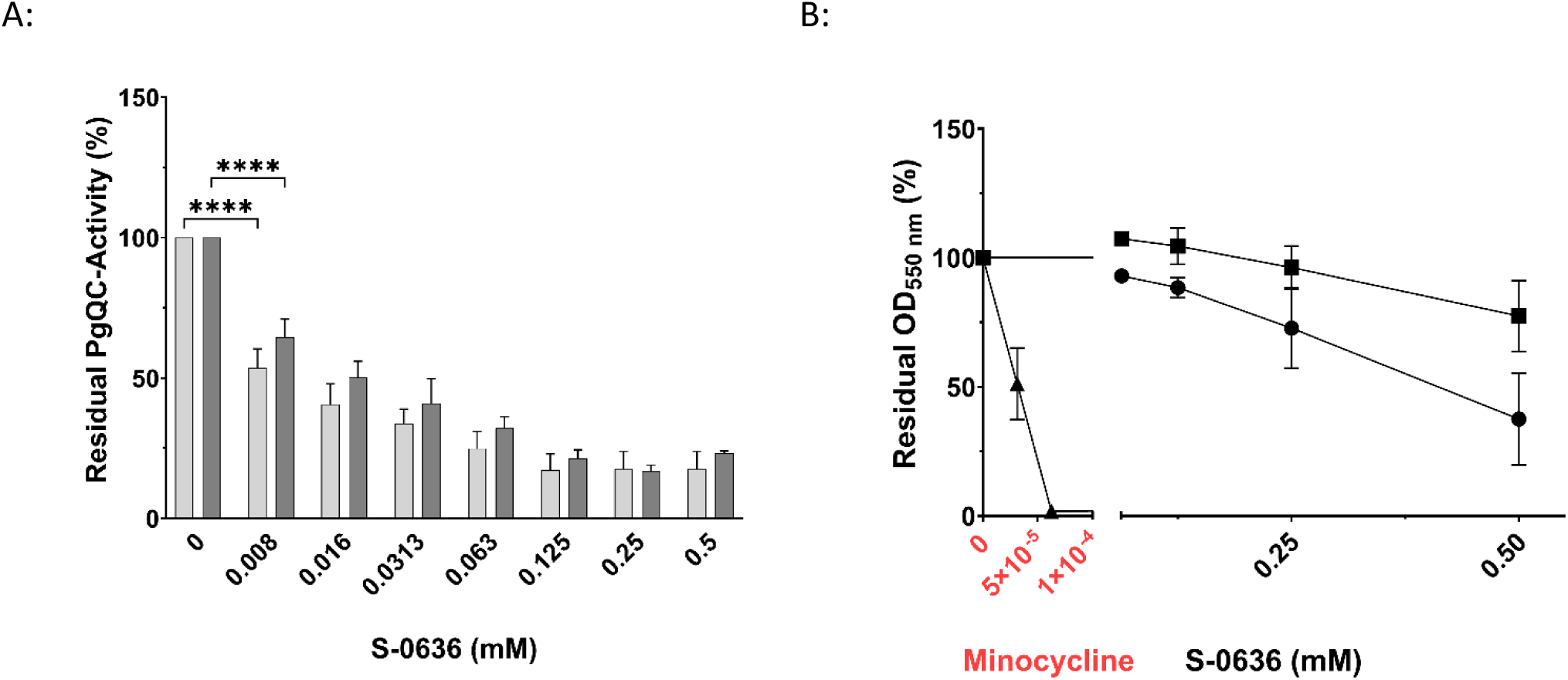
Exposure of bacterial cultures to S-0636 results in a reduction of PgQC activity (A) while maintaining bacterial viability. **(B)** Overnight cultures of *P. gingivalis* ATCC 33277 (A: dark grey bars, B: ■) or *P. gingivalis* W83 (A: light grey bars, B: ●) were diluted and further incubated with a serial 1:2 dilution of S-0636 at 37°C under anaerobic conditions. For comparison, *P. gingivalis* ATCC 33277 was also cultured in the presence of the antibiotic minocycline (▲). After 42 hours, the optical densities were determined, and the samples were adjusted to an OD_600_ of 2 for enzymatic PgQC activity assay. The activity was determined as described above. Experiments were performed in at least three independent technical replicates. Bars denoted by (****) indicate significant difference at p<0.0001 by two-way analysis of variance (ANOVA) with Dunnett’s post hoc test.

To demonstrate that S-0636 functions as a pathoblocker rather than as a classic antibiotic, several *P. gingivalis* strains were treated with either S-0636 or minocycline. Minocycline, a well-known tetracycline derivative, acts as a broad-spectrum antibiotic by inhibiting protein synthesis through binding to the 30S ribosomal subunit in bacteria. It has been used for decades to treat periodontitis. As previously shown, the minimum inhibitory concentration (MIC) for minocycline in the ATCC strain was 0.0625 µM (Inubushi and Liang, 2020). In contrast, no MIC could be determined for S-0636, as even high concentrations affected bacterial growth without causing cell death. Notably, an 8000-fold higher concentration of S-0636 compared to minocycline still showed no bactericidal effect (Fig. 2B).

### Targeting PgQC by S-0636 result in a reduced gingipain activity

Bochtler and colleagues demonstrated that a glutamine-to-arginine substitution at position 24 in the membrane-bound arginine-specific gingipain Rgp A leads to reduced enzymatic activity, particularly when the soluble and secreted form RgpB is deleted. This indicates that RgpA could be a substrate of PgQC. Consequently, inhibition of PgQC could result in decreased Rgp activity and therefore a reduction of a most prevalence virulence factor of *P. gingivalis*. To test this hypothesis, we incubated *P. gingivalis* strains W83 and ATCC with increasing concentrations of S-0636 until they reached the stationary growth phase, and subsequently measured Rgp activity in cell suspensions. As shown in Figure 3, gingipain activity decreased with increasing concentrations of S-0636 in the medium in both *P. gingivalis* strains, reflecting the previously observed reduction in PgQC activity.

**Fig. 3:**
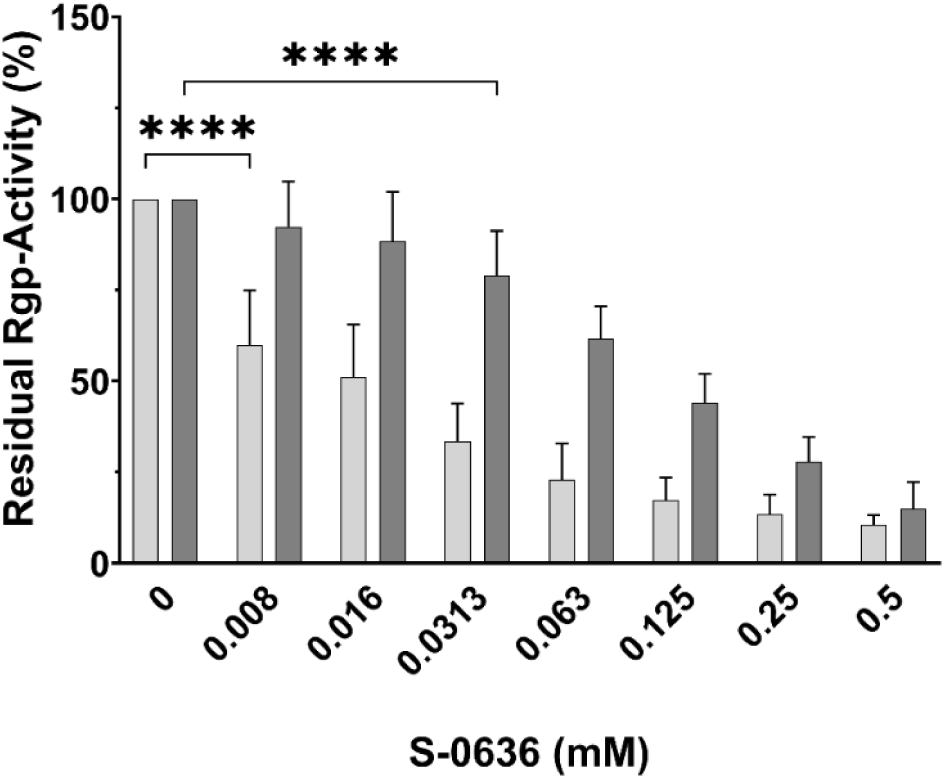
Exposure of bacterial cultures to S-0636 results in a reduction of Rgp activity. Overnight cultures of *P. gingivalis* ATCC 33277 (light grey bars) or *P. gingivalis* W83 (dark grey bars) were diluted and further incubated with a serial 1:2 dilution of S-0636 at 37°C under anaerobic conditions. After 42 hours, the optical densities were determined, and the samples were adjusted to an OD_600_ of 2 for enzymatic Rgp activity as described above. Assay were performed in at least three independent biological experiments. Bars denoted by (****) indicate significant difference at p<0.0001 by two-way analysis of variance (ANOVA) with Dunnett’s post hoc test.

### S-0636 inhibits the *P. gingivalis*-based hemagglutination of red blood cells

*P. gingivalis* possesses the ability to hemagglutinate, i.e., to bind red blood cells, a process that plays a central role in its pathogenesis and survival within the host. Hemagglutination facilitates tissue colonization and ensures access to heme, an essential iron source (Dingsdag et al., 2014; Smalley and Olczak, 2017; Śmiga and Olczak, 2025). The molecular mechanism underlying this process is complex and involves multiple proteins. Key contributors include Rgp A, with its C-terminal adhesion domain, and HagA (Shi et al., 1999; Zhang et al., 2016). Sequence analyses revealed that HagA may also be subject to N-terminal glutamine exposure and thus a potential substrate of PgQC. Consequently, inhibition of PgQC could impair hemagglutination capacity. To investigate this, *P. gingivalis* W83 was incubated with increasing concentrations of the PgQC inhibitor S-0636 for 42 h. Bacterial suspensions were adjusted to OD_600_ of 3 incubated with defibrinated sheep blood at various dilutions. As shown in Figure 4A, *P. gingivalis* exhibited a progressively reduced ability to bind erythrocytes following incubation with higher concentrations of S-0636. Quantitative analysis of hemagglutination across dilution levels confirmed this trend (Figure 4B).

**Fig. 4:**
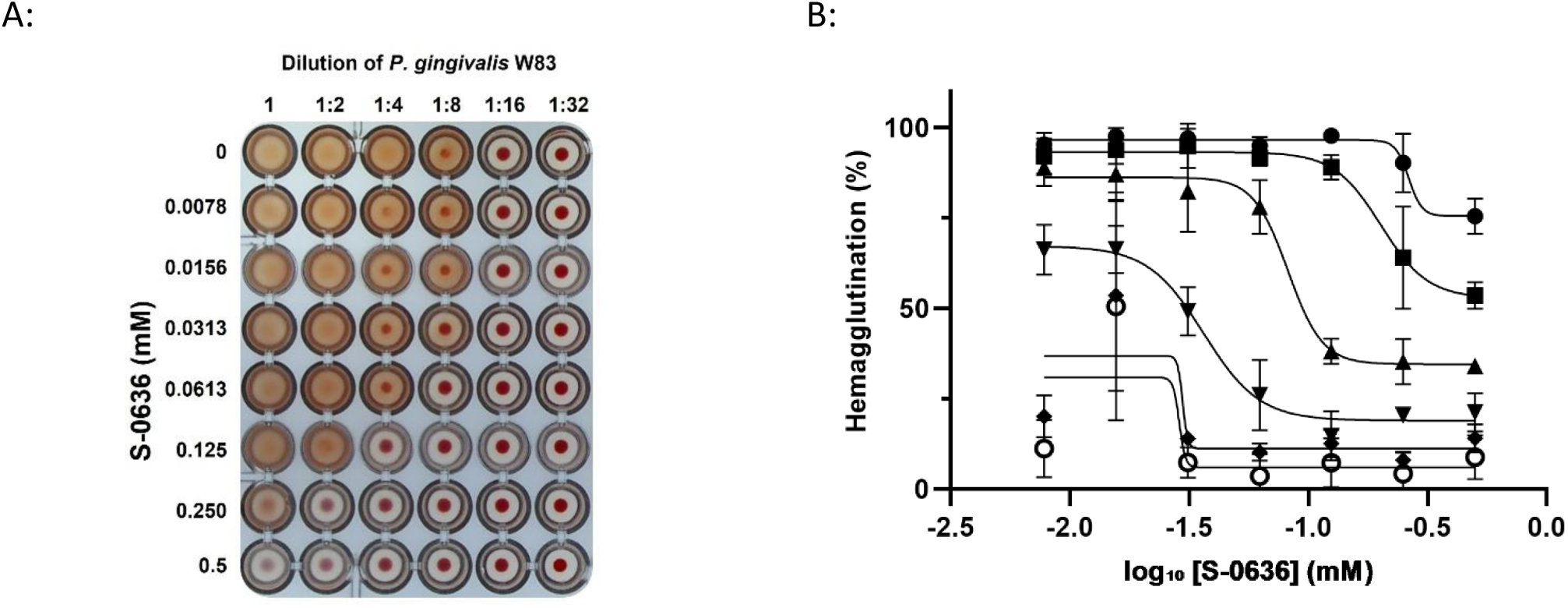
Cultivation of *P. gingivalis* W83 in the presence of S-0636 leads to a decreased ability to mediate hemagglutination. Overnight cultures of *P. gingivalis* W83 were diluted and further incubated with a serial 1:2 dilution of S-0636 at 37°C under anaerobic conditions. After 42 hours, the optical densities were determined, and the samples were adjusted to an OD_600_ of 3. Resulting *P. gingivalis* W83 samples were undiluted (●) or 1:2 (■), 1:4 (▲), 1:8 (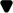), 1:16 (◆) and 1:32 (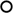) diluted and incubated with a 2% sheep blood solution as described above. The hemagglutination capacity was determined both visually (A) and quantitatively (B), by using the program ImageJ to determine signal intensities and fitting the corresponding curves with GraphPad Prism. Hemagglutination assay was performed in at least three independent biological experiments.

### S-0636 effectively inhibits the invasion of keratinocytes by *P. gingivalis*

The infection process of *P. gingivalis* into epithelial cells involves a sophisticated sequence of molecular interactions and host cell manipulations. The first step is adhesion to epithelial cells. It has been shown that *P. gingivalis* strain ATCC uses its major fimbriae (FimA) to bind to α5β1-integrin receptors on epithelial cells, followed by endocytosis of the bacterium (Yilmaz et al., 2003, 2002).

Since *P. gingivalis* strain W83 is considered a non-fimbriated phenotype and possesses only a low amount of FimA, other virulence factors must play a crucial role in the infection process. Indeed, gingipains, with their adhesion domains, and other adhesins such as HagA, are essential for bacterial adherence (Chen et al., 2001). Moreover, outer membrane vesicles (OMVs) secreted by *P. gingivalis* actively support its invasion into epithelial cells (Farrugia et al., 2020; Ho et al., 2015). These OMVs are rich in virulence factors such as gingipains, fimbriae, and lipopolysaccharides (LPS), which help to manipulate host cells (Okamura et al., 2021).

OMVs are internalized by epithelial cells via endocytosis, and this uptake occurs more efficiently than with whole bacteria. Once inside, OMVs disrupt cellular functions by degrading epithelial proteins, thereby facilitating cell adhesion and migration (Ho et al., 2015). Thus, OMVs play a central role in enhancing bacterial entry, intracellular survival, and immune evasion during the infection process. To investigate whether the application of compound S-0636 reduces the virulence factor activity and thereby the infectivity of *P. gingivalis* W83, the bacteria were incubated with increasing concentrations of S-0636 for 42 hours. Subsequently, gingival keratinocytes were infected with a multiplicity of infection (MOI) of 200 for 90 minutes. After infection, the keratinocytes were lysed, and the lysates were serially diluted and plated on blood agar to determine the colony-forming units (CFU) of *P. gingivalis*. As shown in Figure 5, increasing concentrations of S-0636 led to a progressive reduction in bacterial infectivity. Notably, even a relatively low concentration of 62.5 µM S-0636 reduced the infectivity of *P. gingivalis* W83 by 76%.

**Fig. 5:**
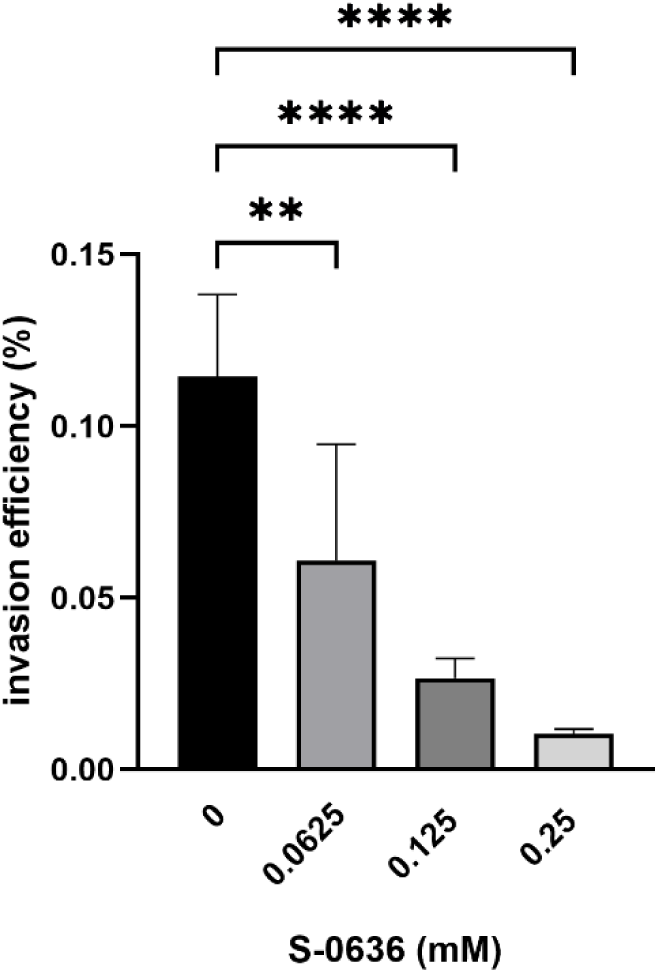
S-0636-mediated reduction of *P. gingivalis* W83 invasiveness. Overnight cultures of *P. gingivalis* W83 were diluted and further incubated with a serial 1:2 dilution of S-0636 at 37°C under anaerobic conditions. After 42 hours, the optical densities were determined, and the samples were adjusted to an MOI of 200 in keratinocyte cell culture medium, followed by a 90-min infection with hTERT TGKs. After cell lysis of TIGKs, different dilutions were plated on blood agar plates to determine the CFU of invasive *P. gingivalis* W83 and incubated for 5-7 days at 37°C under anaerobic conditions. An Infection assay was performed in at least three independent biological experiments. Bars denoted by (**) and (****) indicate significant difference at p= 0.0021 and p<0.0001 by one-way analysis of variance (ANOVA) with Dunnett’s post hoc test.

### S-0636 does not harm other commensal bacteria in the microbiome

The oral cavity hosts the second largest and most diverse microbiota in the human body after the gut, comprising over 700 bacterial species (Caselli et al., 2020; Deo and Deshmukh, 2019). The oral microbiome represents a highly diverse and dynamic microbial ecosystem that plays a fundamental role in maintaining both oral and systemic health. One of its key functions is colonization resistance, where commensal microorganisms occupy ecological niches and prevent the establishment and proliferation of pathogenic species. This is achieved through competitive exclusion, nutrient limitation, and the production of antimicrobial compounds such as bacteriocins and organic acids. The oral microbiome also contributes to oral homeostasis by regulating local pH, supporting epithelial barrier integrity, and facilitating tissue repair. Moreover, it plays a critical role in modulating the host immune system, promoting immune tolerance, and preventing excessive inflammatory responses that could lead to tissue damage (Deo and Deshmukh, 2019). In addition, oral microbes are involved in nutrient metabolism, including the breakdown of dietary carbohydrates and proteins, and the production of bioactive metabolites such as short-chain fatty acids, which influence both microbial and host cell functions. In summary, the oral microbiome performs several essential functions in maintaining oral and overall health and should not be negatively affected.

To rule out an inhibitory effect of S-0636 on bacteria representing a healthy oral microbiome, ten commensal strains were selected based on literature evidence (Abusleme et al., 2021), and their growth was evaluated in the presence of increasing concentrations of S-0636. As shown in Figure 6, none of the tested core strains exhibited growth inhibition at concentrations up to 0.25 mM. At concentrations exceeding this threshold, modest growth inhibition was observed for *V. parvula*, which may reflect nonspecific toxic effects rather than targeted antimicrobial activity.

**Fig. 6:**
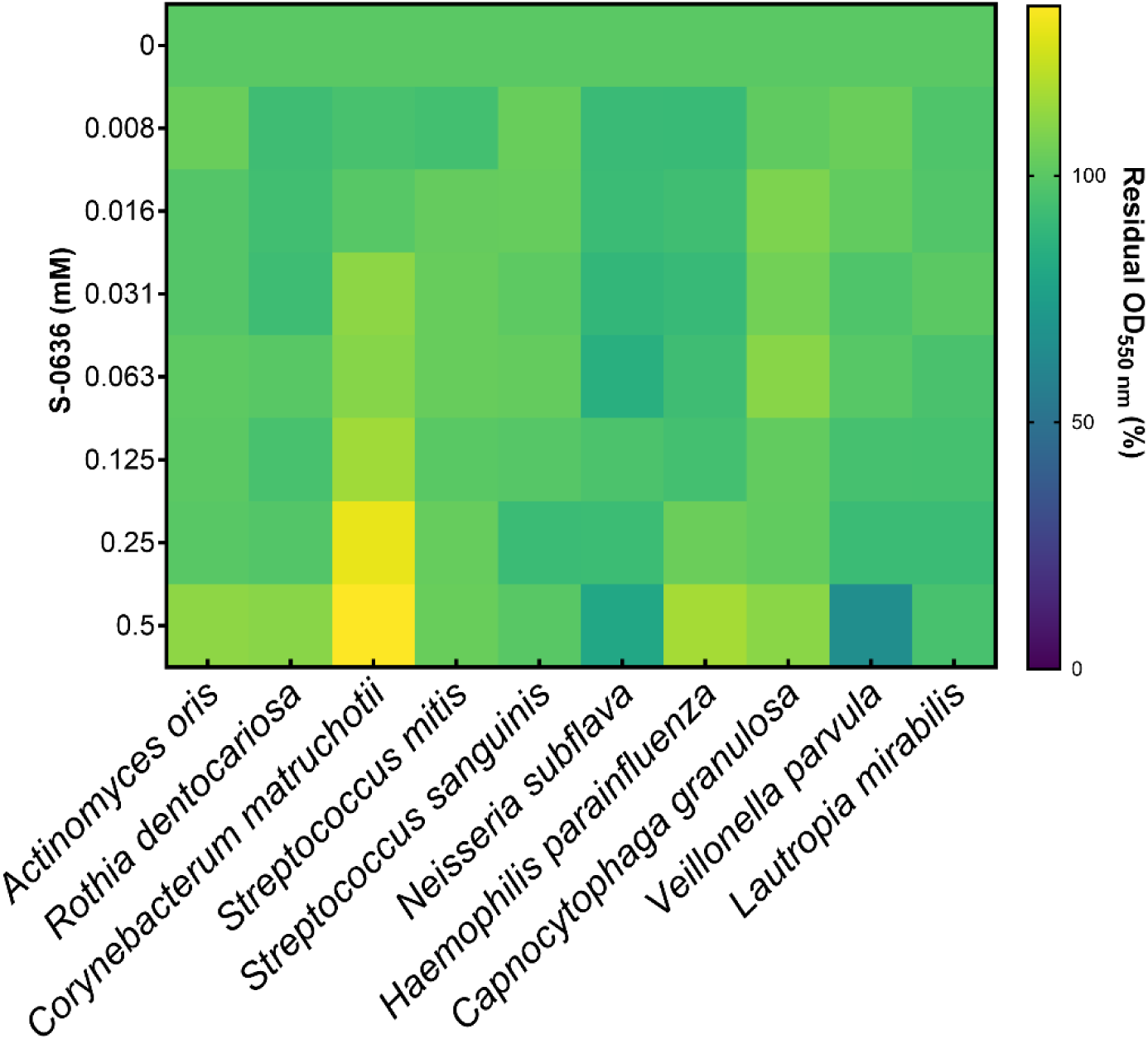
S-0636 does not affects the growth of most oral commensals. Overnight cultures of different bacterial strains were diluted and further incubated with a serial 1:2 dilution of S-0636 under specific conditions as described in the supplementary information. After a certain period, the optical density at 550 nm was measured, and the residual growth rate was determined using the culture w/o S-0636 as a reference. The results were visualized as a heat map using GraphPad Prism10. Growth experiments were performed in at least three independent biological experiments.

### S-0636 induces no resistance mechanism in *P. gingivalis*

Due to selection pressure, antibiotics can induce resistance mechanisms in bacteria, allowing them to evade bactericidal effects. Standard resistance testing methods are often employed to evaluate such potential effects. Typically, bacteria are cultured over multiple passages in the presence of sub-inhibitory concentrations, commonly ¼ of the minimum inhibitory concentration (MIC), followed by reassessment of the MIC for the compound in question (Kowalska-Krochmal and Dudek-Wicher, 2021).

In contrast, pathoblockers such as S-0636 do not exhibit bactericidal activity, and therefore, MIC-based approaches are not suitable and require modification (Filipić et al., 2024; Totsika, 2016). To nonetheless investigate whether S-0636 imposes selection pressure on *P. gingivalis* that could lead to the development of resistance mechanisms, various *P. gingivalis* strains were cultured in the presence of 16 µM S-0636 through 50 successive passages.

After every tenth passage, the strains were cultured without S-0636, and subsequently incubated for 42 hours with increasing concentrations of S-0636. The Rgp activity, serving as a functional readout for the effect of S-0636, was then quantified. As shown in Figure 7, continuous cultivation of *P. gingivalis* in the presence of 16 µM S-0636 over 50 passages did not result in a reduction of gingipain activity, nor did it alter its functional profile. These findings suggest that prolonged exposure to S-0636 does not induce resistance mechanisms in *P. gingivalis*.

**Fig. 7:**
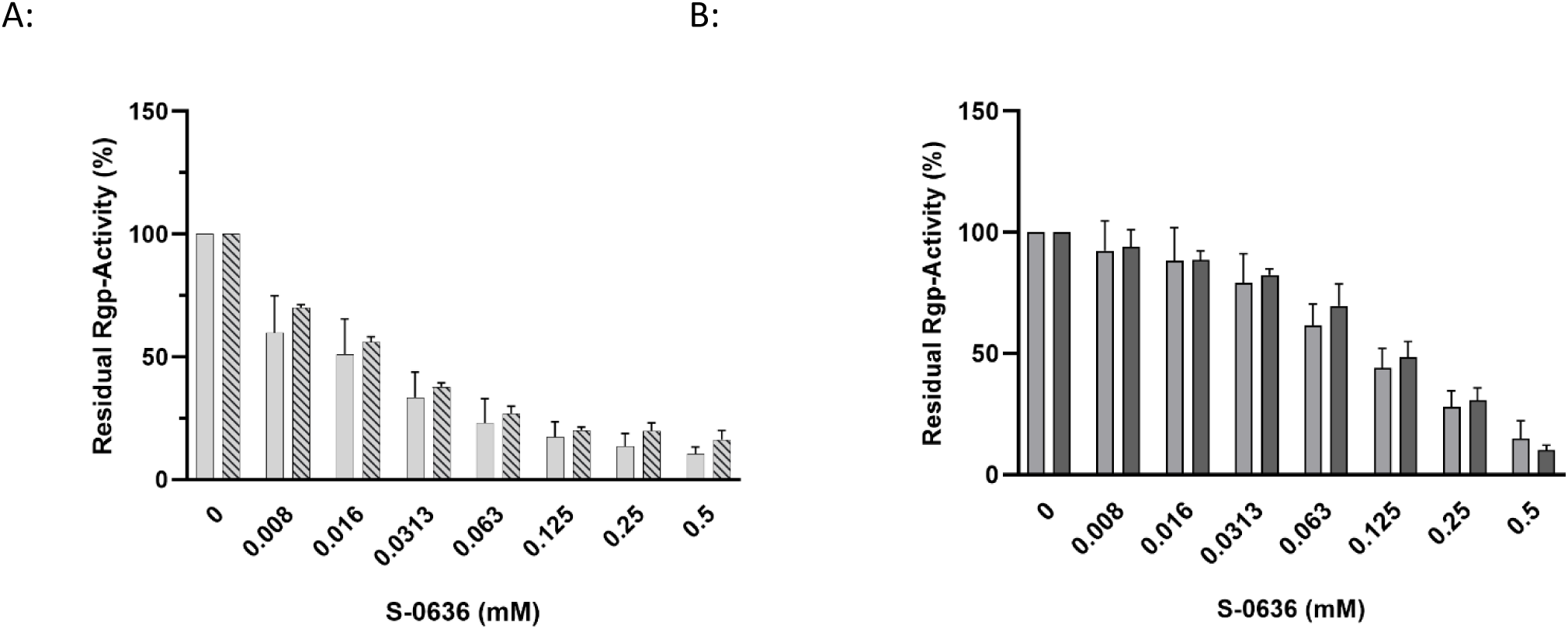
S-0636 exhibits a low potential for resistance development of *P. gingivalis*. *P. gingivalis* ATCC 33277 (A) and W83 (B) were passaged on blood-agar plates containing w/o S-0636 and 16 µM S-0636 and incubated under anaerobic conditions at 37°C for up to 4 days. After every tenth passage, the strains were cultured overnight in the absence of the compound, followed by an incubation with different concentrations of S-0636. After 42 hours, the optical densities were determined, and the samples were adjusted to an OD_600_ of 2 for the enzymatic Rgp activity assay as described above. The striped bars represent the Rgp activity of *P. gingivalis* strains ATCC 33277 and W83 after 50 passages on agar containing S-0636. Assays were performed in at least three independent biological experiments.

### S-0636 demonstrates minimal cytotoxic effects, indicating its potential safety for host cells

In the context of a potential application of S-0636 in oral care, it is essential during the preclinical phase to evaluate the compound’s cytotoxicity toward eukaryotic cells. Therefore, various concentrations of S-0636 were incubated for 24 hours with different cell lines, including gingival keratinocytes, gingival fibroblasts, embryonic kidney cells, and hepatocytes. Following incubation, cell morphology and viability were assessed using the WST-8 assay, which is based on the enzymatic reduction of the tetrazolium salt WST-8 to a soluble formazan product by mitochondrial and cytosolic dehydrogenases in metabolically active cells. This reaction enables a quantitative determination of cell viability via spectrophotometric measurement. As shown in Figure 8 gingival keratinocytes, as well as kidney and liver cells, maintained a viability above 80% after 24 hours of exposure to 0.5 mM S-0636, whereas gingival fibroblasts exhibited a viability of approximately 60%. These observations suggest that non-specific effects may contribute to the reduced metabolic activity observed in fibroblasts. The exact mechanisms of these effects are currently under investigation.

**Fig. 8:**
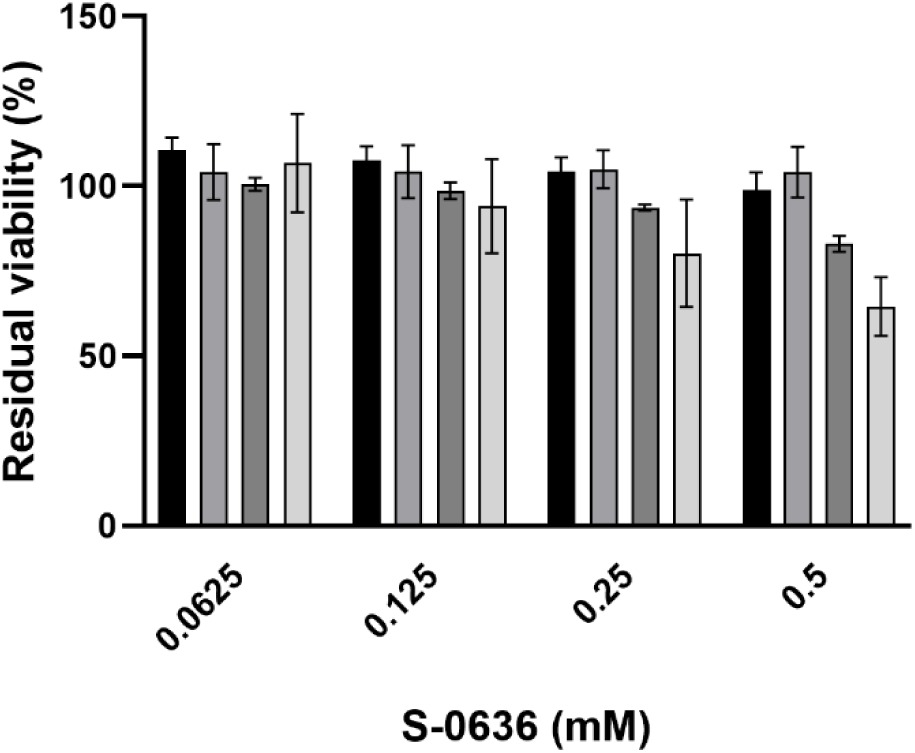
S-0636 exhibits a low potential of Cytotoxicity in human cell lines. Human liver cells (HepG2, black bars), human embryonic kidney cells (HEK293, grey bars), human gingival keratinocytes (hTERT-TIGK, dark grey bars), and human gingival fibroblasts (hTERT-TIGF, light gray bars) were seeded and cultivated as described above. Once the cells reached ∼80% confluency, the medium was replaced with fresh medium containing various concentrations of S-0636, followed by an additional 24-hour incubation under the same conditions. The WST-8 assay determined the viability of cells. Cells treated with 0.1% Triton X-100 served as a negative control), while cells treated with the S-0636 compound diluent served as a positive control. Cell viability was quantified by measuring the absorbance of WST-8 formazan at 450 nm using a Tecan Sunrise microplate reader. All experiments were performed in at least three independent biological replicates.

## Discussion

Severe periodontitis affects about one billion people worldwide (12.5% prevalence), ranking among the six most common human diseases (James et al., 2018; Nascimento et al., 2024; Wu et al., 2025). Only untreated dental caries surpasses it, with over three billion incident cases and two billion prevalent cases annually (James et al., 2018). Compared with other oral infections, periodontitis remains the leading chronic inflammatory oral disorder, underscoring a public-health burden that rivals or exceeds many systemic infectious diseases (Hu et al., 2025; Nascimento et al., 2024). Besides manual debridement, existing approaches have their limitations. Therefore, chlorhexidine remains the gold-standard antiseptic adjunct in periodontitis, primarily due to its high substantivity, which sustains bacteriostatic levels in crevicular fluid for approximately 12-24 hours after a single rinse (Deus and Ouanounou, 2022; Thangavelu et al., 2020). In contrast, a meta-analysis shows only approximately 0.12 mm additional reduction in probing depth over six months when CHX-gel is added to scaling and root planning (Zhao et al., 2020). In addition, long-term daily use (≥ 4 weeks) triggers marked extrinsic staining, taste alteration, and mucosal irritation, while fostering microbiome shifts toward cariogenic streptococci and the enrichment of tetracycline efflux resistance genes (Deus and Ouanounou, 2022). Accordingly, extended prescriptions are typically reserved for short postsurgical or refractory phases. Although in most of the cases an effective option, systemic antibiotics foster antimicrobial resistance, may cause systemic side-effects including allergies and gastrointestinal issues, can interact with common medications, are known to disrupt gut and oral microbiota homeostasis, and often fail to reach bactericidal levels inside biofilm-protected pockets (Abe et al., 2022; Barca et al., 2015; Heitz-Mayfield, 2009; Heta and Robo, 2018; Slots, 2004; Yuan et al., 2023). In contrast, locally delivered antibiotics have the advantage of a limited systemic exposure but the penetration is often inconsistent due to complex root anatomy (Abu-Ta’a and Bazzar, 2023; Ilyes et al., 2024; Yusri et al., 2021). They may release cytotoxic concentrations of the compound and can provoke mild gingival reactions (Matesanz-Pérez et al., 2013). In contrast they yield only marginal additional clinical attachment gains of ca. 0.3 - 0.7 mm and add cost and logistical burden (Yusri et al., 2021). Consequently, current periodontal guidelines reserve antibiotics for severe or refractory cases after thorough mechanical debridement (Sanz et al., 2020b).

Recent periodontitis therapy has undergone a paradigm shift towards targeted, drug-based strategies that encompass a broad spectrum of innovative approaches. These include sophisticated nanoparticle delivery systems for site-specific drug targeting (India et al., 2024; Nakajima et al., 2025), antimicrobial and host-modulating peptides that address both pathogen elimination and immune response regulation (Zhang et al., 2024), and specialized pro-resolving mediators such as lipoxin A4 and resolvin E1 that actively promote inflammation resolution (Alshibani, 2022; Dyke, 2017; Hasturk et al., 2021). Additionally, bone anabolic agents such as romosozumab and teriparatide, offer genuine regenerative potential (Cecoro et al., 2021; Reghunath et al., 2024), while immunomodulatory approaches utilizing stem cell products provide novel therapeutic avenues (Pahade et al., 2023; Zhao et al., 2025). Remarkably, most of these innovative approaches are being developed by universities, public institutions, or small biotechnology companies, with virtually only bone anabolic agent development being pursued by major pharmaceutical corporations.

Having the fundamental fact in mind, that periodontitis is a disease with an underlying dysbiosis of the oral microbiome, the question of modulating approaches arises increasingly. Especially after a successful standard-of-care therapy, one major goal is to disrupt the vicious circle of re-colonization of the affected sites with pathogens and allow commensals to occupy the niches. Several approaches were developed to support the microbiome during the supportive periodontitis therapy period between two recalls. Probiotics and prebiotics have emerged as promising adjunctive treatments for periodontitis, offering several therapeutic benefits while presenting certain limitations that must also be considered. Probiotics seem to demonstrate multiple beneficial mechanisms in periodontal treatment. These beneficial microorganisms compete with pathogenic bacteria for adhesion sites on oral mucosa and tooth surfaces and can inhibit the colonization of pathogens such as *P. gingivalis* and *F. nucleatum* (Doucette et al., 2024; Singh et al., 2017). Prebiotics enhance the effectiveness of probiotics by selectively promoting the growth of beneficial microorganisms in the oral cavity. This synergistic effect helps to re-establish the microbiome balance (Paradowski et al., 2024). Despite promising results, several limitations constrain the widespread adoption of probiotics in periodontal therapy. Clinical benefits are often short-term, and the diversity of probiotic strains, delivery methods, dosages, and treatment durations across several studies makes it difficult to establish standardized treatment protocols (Jayaram et al., 2016; Schlagenhauf and Jockel-Schneider, 2021). Interestingly, different meta-analyses yield conflicting results, ranging from “no additional benefit for pocket reduction” to “limited, short-term benefits in attachment gain” (Schlagenhauf and Jockel-Schneider, 2021). Consequently, the European Association of Periodontology currently does not recommend probiotics as adjunctive therapy to subgingival instrumentation due to insufficient evidence

A highly promising strategy in this context is the incorporation of pathoblockers into oral care formulations, representing a novel application within the framework of next-generation antimicrobials (NGAs). This approach offers several distinct advantages. First, bacteria are not killed, which significantly reduces the potential risk of the development resistance mechanism. Secondly, the targeted inhibition of virulence in keystone pathogens preserves the integrity of the remaining oral microbiota, leaving commensal species unaffected. This selection pressure relief enables beneficial microorganisms to reoccupy ecological niches that pathogenic species would otherwise dominate during the progression of disease. Unlike probiotic approaches, which rely on the introduction of exogenous microorganisms, this strategy promotes microbiome restoration and stabilization through the host’s endogenous microbial community. From a pharmaceutical and formulation standpoint, the use of a chemically defined small molecule offers distinct advantages in terms of stability, scalability, and regulatory feasibility compared to oral care products containing live microorganisms or peptide-based agents. The pathoblocker approach, discussed here, was pioneered through the discovery of bacterial type II glutaminyl cyclase, which is conserved among bacteria of the phylum Bacteroidetes (Bochtler et al., 2018; Taudte et al., 2021). This enzyme catalyzes the cyclization of N-terminal glutamine residues in a wide range of proteins, representing an essential post-translational modification of many virulence factors in P. gingivalis. If this cyclization does not occur, it affects the functionality of these proteins, either through reduced stability, impaired secretion, or incomplete maturation (Bochtler et al., 2018; Szczęśniak et al., 2023). The detailed mechanism underlying this modification remains the subject of ongoing research. Nevertheless, it is undisputed that bacterial glutaminyl cyclase type II represents an attractive target for the development of selective small molecules, which could potentially be used as pathoblockers to support therapeutic interventions (Ramsbeck et al., 2021; Taudte et al., 2021). We therefore further investigated active compounds and used one promising candidate for the target validation, which is reported here, compound S-0636. It is a new chemical entity with a high activity against PgQC combined with a promising profile regarding physico-chemical properties. So, it was used for a further evaluation of the hypothesis of a possible selective inhibition of *P. gingivalis* virulence.

As an initial step, it was essential to demonstrate that S-0636 not only inhibits the isolated enzyme but also exhibits inhibitory activity under physiological conditions in planktonic *P. gingivalis* cells. The observed inhibition of QC activity provides compelling evidence that the compound effectively penetrated into the periplasm, in both the non-encapsulated *P. gingivalis* ATCC strain and the encapsulated W83 strain. This penetration may be attributed to the compound’s high polarity and aqueous solubility. Several publications have analysed compounds according to their efficacy against Gram-negative bacteria, revealing a preference for smaller, polar, and water-soluble compounds. It is hypothesized that these properties facilitate the transport through bacterial aquaporins (Ebejer et al., 2016; Mugumbate and Overington, 2015; O’Shea and Moser, 2008). S-0636, the compound employed in this study, combines these favourable physicochemical features within its chemical structure. With its molweight of 325 Da, a topological polar surface area (TPSA) of 124.4 Å^2^ (Ertl et al., 2000) and measured logD values between −2.01 (logD_7.0_) and −1.68 (logD_9.0_) (data not presented) it harbours the described required physio-chemical properties to act on gram-negative bacteria. Even in the slight basic pH-range around pH = 8.0, where *P. gingivalis* is unperturbed and the PgQC activity is optimal (Taudte et al., 2021), the guanidinium group of S-0636 is predicted to be ionized.

To further substantiate our hypothesis, we next aimed to demonstrate that inhibition of glutaminyl cyclase (PgQC) by S-0636 leads to a measurable reduction in the virulence of *P. gingivalis*. Previous studies by Bochtler (Bochtler et al., 2018), and subsequently by Szczesniak (Szczęśniak et al., 2023) have shown that more than 70% of all secreted proteins in *P. gingivalis* are likely substrates of PgQC. Sequence analysis of key virulence factors (Silva and Cascales, 2021) using Signal P revealed that several components of the Type IX secretion system, such as PorV and PorN (KEGG Genome entries PGN_0023 and PGN_1673) as well as specific proteases including the arginine gingipains RgpA (PGN_1970) and RgpB (PGN_1466), dipeptidyl peptidase IV (DPP IV, PGN_1469), peptidylarginine deiminase (PPAD, PGN_0898) and prolyl tripeptidyl peptidase A (PTP A, PGN_1149) are putative QC substrates. In addition, adhesion proteins such as HagA (PGN_1733) and the heme-binding protein (PGN_0659), along with proteins essential for the biogenesis and release of outer membrane vesicles (OMVs), are also predicted to undergo QC-mediated N-terminal modification. Our results indicate that inhibition of PgQC has a direct impact on the released activity of virulence factors. Hence, we demonstrated that bacterial pre-incubation with S-0636 attenuates the Rgp-gingipain response in both *P. gingivalis* strains. Moreover, upstream proteases such as DPP IV and PTP A are also targeted, suggesting a broader modulatory effect on the proteolytic cascade (data not shown). As previously described proteases, especially gingipains, play a central role in the pathogenicity and survival of *P. gingivalis*. As an asaccharolytic organism, *P. gingivalis* relies on proteolysis to obtain amino acids from host proteins as its primary energy source (Nemoto and Ohara-Nemoto, 2016). In addition, they promote biofilm formation, facilitate host colonization and invasion, degrade host proteins such as immunoglobulins and complement components, and contribute to immune evasion and chronic inflammation (Hočevar et al., 2020). In summary, reduced protease activity profoundly impairs the overall fitness and pathogenic potential of *P. gingivalis* (Pedrosa et al., 2025). The coordinated action of multiple virulence factors often leads to optimal pathogenic outcomes. For instance, not only does membrane-associated RgpA mediate host cell invasion, but additional proteins containing adhesion domains such as HagA and the hemin-binding proteins also contribute to adhesion to erythrocytes and other host structures. The ability to bind erythrocytes, referred to as hemagglutination, can be assessed using a hemagglutination assay. To investigate whether this ability is affected by S-0636, various *P. gingivalis* strains were pre-incubated with increasing concentrations of S-0636 and subsequently incubated with defibrinated sheep blood. We observed that pre-incubation with S-0636 correlated with a reduced capacity to bind erythrocytes. Furthermore, different *P. gingivalis* strains and clinical isolates, such as M5-1-2 and R14, exhibited varying degrees of erythrocyte binding. However, in all cases, growth in the presence of S-0636 resulted in decreased erythrocyte adhesion (data not shown).

Adhesion to host cells and disruption of tissue structural integrity represents the initial steps in the infection process of *P. gingivalis*. This is followed by transcytosis, which typically involves three stages: entry (via endocytosis), intracellular survival, and exit (Jongh et al., 2023). A key difference between the *P. gingivalis* strains ATCC 33277 and W83 lies in their surface structures: ATCC 33277 expresses surface-associated long fimbriae of type I, whereas W83 possesses a polysaccharide capsule of serotype K1. Consequently, W83 is classified as a non-fimbriated, encapsulated type. This capsule confers increased resistance to host-mediated phagocytosis, enhances bacterial survival in hostile environments, and leads to a reduction in the secretion of pro-inflammatory cytokines by the host (Płonczyńska et al., 2025). Additionally, W83 exhibits higher gingipain activity compared to other strains (Reyes et al., 2013). These combined features contribute to the elevated virulence and invasiveness of the W83 strain (Murugaiyan et al., 2024; Rodrigues et al., 2012). For this reason, *P. gingivalis* W83 was selected for our infection experiments to demonstrate that the compound attenuates the pathogenicity of the encapsulated type of *P. gingivalis*. Our data demonstrate that the growth of *P. gingivalis* W83 in the presence of S-0636 results in a marked reduction in bacterial invasiveness. Notably, pre-incubation with only 0.0625 mM of the compound significantly impairs the ability of *P. gingivalis* to infect oral keratinocytes. Which specific step of the infection process is affected by the inhibition of PgQC remains unclear. However, it is highly likely that the observed reduction in invasiveness results from the combined impairment of multiple factors, especially considering that several virulence determinants are known substrates of PgQC.

In conclusion, our previous findings indicate that inhibition of PgQC by compound S-0636 results in a significant reduction of gingipain activity, impairs the hemagglutination capacity of *P*. *gingivalis*, and diminishes its ability to infect gingival keratinocytes. Given that S-0636 is a highly specific small-molecule inhibitor and PgQC is conserved within the phylum Bacteroidetes. Nevertheless, to exclude potential non-specific effects, particularly on commensal members of the healthy oral microbiota (Abusleme et al., 2021), representative bacterial strains were selected and incubated with increasing concentrations of compound S-0636. We demonstrated that the growth of ten core members of a healthy biofilm remains unaffected following the application of the compound. This finding again reflects the selectivity of the approach; however, it is worth noting that these ten strains do not represent the complete picture of the several hundred different bacterial species that comprise the oral microbiome. Currently, it cannot be ruled out that S-0636 exerts nonspecific effects against other commensal bacteria.

A key advantage of our strategy is the low potential of S-0636 to induce resistance mechanisms in *P. gingivalis*. Although the compound is targeting PgQC, an essential enzyme whose genetic deletion results in a lethal phenotype, our approach is based on competitive inhibition that permits residual enzymatic activity. This minimal activity appears sufficient to sustain bacterial viability, while substantially impairing virulence (Szczęśniak et al., 2023).

In contrast to conventional antibiotics, such as those used in the van-Winkelhoff-Cocktail (Amoxicillin and Metronidazole), which exert strong bactericidal effect and thereby promote the development of resistance mechanism, our approach selectively attenuates pathogenicity without compromising bacterial survival. This virulence-targeted strategy reduces selection pressure and offers a promising avenue for the development of next-generation antimicrobials (NGAs) with improved safety and resistance profiles. To evaluate this theoretical advantage, we conducted a classical resistance development assay in which *P. gingivalis* was exposed to S-0636 at sub-inhibitory concentrations over 50 consecutive passages. Following 20, 30, 40, and 50 passages, the sustained efficacy of S-0636 was assessed by measuring its inhibitory effect on Rgp release. Remarkably, no diminution in compound responsiveness was observed throughout the entire passage series, suggesting an absence of resistance development under these experimental conditions. These findings must be interpreted with appropriate caution, as conventional MIC-based resistance testing methodologies are fundamentally incompatible with pathoblocker evaluation principles. Traditional resistance assays are designed to detect changes in bacterial survival or growth inhibition, parameters that are irrelevant for compounds that specifically target virulence mechanisms without affecting bacterial viability. In the absence of established alternative methodologies for assessing pathoblocker resistance, we employed this classical experimental framework to obtain preliminary insights into potential resistance development patterns. While these results provide encouraging initial evidence supporting the theoretical resistance-reduction benefits of the pathoblocker approach, the development of specialized resistance testing protocols tailored to anti-virulence compounds remains an important area for future methodological advancement.

In general, it should be noted that these results were generated exclusively with planktonic bacterial cultures. It is well established that effects observed on isolated bacterial species often differ significantly or may be absent entirely in mixed biofilms or complex microbiomes (Bengtsson-Palme, 2020; Sadiq et al., 2023). In bacterial communities, the loss of functions in one species may be compensated by the activity of other biofilm members. Additionally, the extracellular matrix plays a crucial role, supporting different strains and shielding them from external influences.

Future experiments will focus on evaluating S-0636’s effects on more complex biofilms and investigating its indirect modulation of immunocompetent cells. Despite its experimental and technical limitations, the results presented here demonstrate a highly promising novel approach for modulating and stabilizing a healthy oral microbiome, with further investigations currently in progress.

## Author Contributions

Conceptualization: NT and MB, experiments: NJ, LL, NT, DR, writing: NT and MB, editing and revision: JP, SE, SS and DR, resources/funding: MB and SE, coordination: NT and MB

## Acknowledgments

We gratefully thank D. Meitzner and S. Vogt for bio-analytical and S. König for synthetic support.

## Data Availability Statement

The raw data supporting the conclusions of this article will be made available by the authors, without undue reservation.

